# Host transcriptomic profiling of CD-1 outbred mice with severe clinical outcomes following infection with *Orientia tsutsugamushi*

**DOI:** 10.1101/2022.05.05.490713

**Authors:** Joseph Thiriot, Yuejin Liang, James Fisher, David H. Walker, Lynn Soong

## Abstract

*Orientia tsutsugamushi* is an obligately intracellular bacterium with endothelial tropism and can cause mild to lethal scrub typhus in humans. No vaccine is available for this reemerging and severely neglected infection. Previous scrub typhus studies have utilized inbred mice, yet such models have intrinsic limitations. Thus, the development of suitable mouse models that better mimic human diseases is in great need for immunologic investigation and future vaccine studies. This study is aimed at establishing scrub typhus in outbred CD-1 mice and defining immune biomarkers related to disease severity. CD-1 mice received *O. tsutsugamushi* Karp strain via the i.v. route; major organs were harvested at 2-12 days post-infection for kinetic analyses. We found that for our given infection doses, CD-1 mice were significantly more susceptible (90-100% lethal) than were inbred C57BL/6 mice (0-10% lethal). Gross pathology of infected CD-1 mouse organs revealed features that mimicked human scrub typhus, including pulmonary edema, interstitial pneumonia, perivascular lymphocytic infiltrates, and vasculitis. Alteration in angiopoietin/receptor expression in inflamed lungs implied endothelial dysfunction. Lung immune gene profiling using NanoString analysis displayed a Th1/CD8-skewed, but Th2 repressed profile, including novel biomarkers not previously investigated in other scrub typhus models. Bio-plex analysis revealed a robust inflammatory response in CD-1 mice as evidenced by increased serum cytokine and chemokine levels, correlating with immune cell recruitment during the severe stages of the disease. This study provides an important framework indicating a value of CD-1 mice for delineating host susceptibility to *O. tsutsugamushi*, immune dysregulation, and disease pathogenesis. This preclinical model is particularly useful for future translational and vaccine studies for severe scrub typhus.

**Author Summary:** Scrub typhus is a severely neglected and potentially fatal disease caused by *Orientia tsutsugamushi*, a genetically intractable, obligately intracellular bacterium that annually infects at least one million people worldwide. There is no vaccine available, and our current understanding of the host immunological response and mechanisms remains very limited. Appropriate animal models of infection that recapitulate the disease are essential to the development of effective therapeutics and vaccines. In this study, we characterized the immunologic responses by transcriptomics and Bio-plex assays in outbred CD-1 mice with lethal *O. tsutsugamushi* infection. We found that CD-1 mice were highly susceptible to infection and that the high mortality correlated with a Th1/CD8-skewed, but Th2 repressed, immune profile during the acute phase of disease. This proinflammatory state was further confirmed by elevated cytokine and chemokine levels in the sera. Collectively, this study established CD-1 mice as a practical, preclinical model to define pathogenic mechanisms underlying severe scrub typhus and for future immunologic and translational studies during *O. tsutsugamushi* infection.

## Introduction

Scrub typhus is caused by *O. tsutsugamushi* (*Ot*), a Gram-negative, lipopolysaccharide-negative intracellular bacterium that preferentially replicates within the cytosol of endothelial cells and phagocytes (monocytes, macrophages, and dendritic cells) [1]. This vector-borne pathogen causes at least one million new cases per year, with a population of one billion at risk of infection [1,2]. While previous studies have indicated this infection is confined to a large area of Southeast Asia known as the ‘tsutsugamushi triangle’, the increasing recognition of cases in other geographical locations such as Africa and South America heighten the need to address this neglected tropical pathogen [1–3]. Effective treatment consists of early use of antibiotics, such as doxycycline, but delayed or missed diagnosis can result in up to 30% fatality rate [4]. Following exposure to chigger bites, some individuals exhibit an eschar at the inoculation site, and other early signs include fever and flu-like symptoms [4]. While many of these cases are self-limiting, patients with severe scrub typhus can develop interstitial pneumonia, hepatic inflammation, and meningoencephalitis [5]. These lesions may lead to acute lung damage, including edema, hemorrhage, diffuse alveolar damage, and interstitial cellular infiltration [6]. This acute tissue damage can lead to multi-organ failure, which is the cause of fatalities.

Murine models of *Ot* infection have been used as far back as 1948 for cross-vaccination tests [7], and have utilized the intraperitoneal (i.p.) route for inoculation [8–10]. Since then, the development of more naturally relevant scrub typhus models via the intradermal (i.d., the natural method of entry) or intravenous (i.v., representing hematogenous spread) routes of inoculation in inbred strains of mice have greatly enhanced our understanding of *Ot*-host interactions and disease pathogenesis [11–21]. Studies with inbred C57BL/6 (B6) and BALB/c mice have provided a stable and consistent background to investigate *Ot* virulence and immunological responses. The use of inbred mice is advantageous for many pilot studies that rely on low in-group variabilities. Despite these advantages, there is an ever present need to characterize how genetic diversity influences the host immunity during infection and vaccination studies, in order to reflect human populations. This need has been brought to light in a study by Martin *et al*. indicating that after protection and infection, the memory CD8 T cell pool size and rate of phenotypic progression were highly variable in individual outbred vs. inbred mice [22]. In some infection systems, inbred mice have been found to be intrinsically less resistant to bacterial and viral challenge, causing a confined view of an infection response [23–26]. Outbred mice have historically been used for pharmacological, toxicology, aging, and cancer studies [27]. They are also valuable in therapeutic and vaccine studies for other infectious diseases because of their broader genetic diversities [27].

Outbred CD-1 Swiss Webster (CD-1) mice, for example, have been recently used for vaccination studies against bacterial, viral, and parasitic infections [23–27]. Their use in the context of *Ot* infection remains limited, although some publications have used CD-1 mice to examine bacterial dissemination either via chigger bites [20,28–32] or needle-based i.d. inoculation [19,33], or to compare virulence among different *Ot* strains/isolates following i.p. inoculation [9,19,20]. To date, only one publication has used CD-1 mice for an *Ot* vaccine-related study [34]. For murine models of scrub typhus research, immunological studies have mostly focused on inbred models [35–37]. The wild-type and gene-targeted knockout mice on the B6 background have provided new insights into the magnitude and kinetics of host immune responses at the tissue and cellular levels during infection [17,35,38–41]. However, the CD-1 mouse model has rarely been utilized, and comprehensive immune profiling is still lacking, as the available publications have only examined basic antibody responses, T cell cytokine levels, lymph node sizes, and general pathology [19,29,32,33]. This has led us to investigate CD-1 mouse immune responses, using methods we have established for acute scrub typhus models in B6 mice (i.v. route) [36,38]. Since lethally infected B6 mice show a Th1-skewed, but Th2-suppressed, inflammatory response during late stages of acute disease, we sought to validate if this polarized immune response is specific to *Ot*-infected B6 mice or is a hallmark for lethal scrub typhus across mouse models.

In this study, we evaluate the benefit of using an outbred mouse model of *Ot* infection to explore the immune response profiles and biological hallmarks of severe scrub typhus. We first compared the susceptibility of CD-1 and B6 mice to *Ot* infection and confirmed the hyper-susceptibility of CD-1 mice, as these mice completely succumbed by day 12 post-infection. Bacterial burdens were prevalent throughout major organs, showing pathological lesions resembling that in scrub typhus patients. Secondly, we performed an immunologic differential expression analysis in CD-1 mice during infection and revealed a highly Th1/CD8-skewed immune response during the acute phase of disease. These findings were complemented through quantification of relevant mRNA levels in the tissues via qRT-PCR, as well as proinflammatory cytokines and chemokines in the serum using Bio-Plex assay. To our knowledge, this is the first report of an i.v. CD-1 mouse model of *Ot* infection, allowing for a comprehensive study for biological hallmarks of severe scrub typhus. This study broadens our understanding of scrub typhus pathogenesis and opens new areas of mechanistic investigation.

## Materials and Methods

### Ethics Statement

The University of Texas Medical Branch (UTMB) complies with the USDA Animal Welfare Act (Public Law 89-544), the Health Research Extension Act of 1985 (Public Law 99-158), the Public Health Service Policy on Humane Care and Use of Laboratory Animals, and the NAS Guide for the Care and Use of Laboratory Animals (ISBN-13). UTMB is a registered Research Facility under the Animal Welfare Act. It complies with NIH policy and has current assurance with the Office of Laboratory Animal Welfare. All procedures were approved by the Institutional Biosafety Committee, in accordance with Guidelines for Biosafety in Microbiological and Biomedical Laboratories. Infections were performed following Institutional Animal Care and Use Committee approved protocols (2101001 and 1902006) at UTMB in Galveston, TX.

### Mouse infection and organ collection

Female Swiss Webster CD-1 outbred mice were purchased from Envigo (East Millstone, NJ). Female B6 mice were purchased from Jackson Laboratory (Bar Harbor, ME). Mice were maintained under specific pathogen-free conditions in the same room for 9-10 days and infected at 8-12 weeks of age. Infections were performed in the Galveston National Laboratory ABSL3 facility at UTMB. All tissue processing and analytic procedures were performed in BSL3 or BSL2 laboratories, respectively. All infections were performed using the same bacterial stock of *O. tsutsugamushi* Karp strain prepared from Vero cells, as described in our previous studies [36,40]. Mice were inoculated i.v. with 5.6 ×10^4^ or 4.32 × 10_4_ focus forming units (FFU), or with PBS as a negative control. Mice were monitored daily for body weight, signs of disease, and disease scores. Tissue samples were collected at days 2, 4, 6, 8, 10, and 12 post-infection and inactivated for immediate and subsequent analysis, with mock infected animals serving as controls.

### Bacterial load quantification

Animal tissues were collected and stored in RNA*Later* (Qiagen) at 4°C overnight for inactivation and then placed at -80°C. DNA was extracted using the DNeasy Blood & Tissue Kit (Qiagen). Bacterial loads in organ tissue were quantified via qPCR, as described previously [38]. Bacterial loads were normalized to total nanogram (ng) of DNA per μL of sample. Data are expressed as the copy number of 47-kDa gene per ng of DNA. The copy number for the 47-kDa gene was determined by serial dilution of known concentrations of a control plasmid containing a single-copy insert of the gene.

### Nanostring gene expression profiling

Total RNA was extracted from mock and *Ot*-infected tissues at 4-, 8-, and 12-days post infection (3 samples/group) using the RNeasy Kit (Qiagen). Samples were processed at the Baylor College of Medicine Genomic and RNA Expression Profiling Core (Houston, TX). The Nanostring nCounter platform and the Mouse Immunology Panel were used to quantify transcripts of 561 immune plus 14 housekeeping genes (NanoString Technologies, Seattle, WA). Gene expression was normalized to housekeeping gene expression, and the data were analysed using the nSolver Software Version 4 and Advanced Analysis Version 2.0 (NanoString Technologies), as described in our previous study [42].

### Quantitative reverse transcriptase PCR (qRT-PCR)

Total RNA was extracted from tissues using the RNeasy mini kit (Qiagen) and treated with DNase according to the manufacturer’s protocol (Qiagen). cDNA was synthesized via the iScript cDNA synthesis kit (Bio-Rad Laboratories). Target gene abundance was measured by qRT-PCR using a Bio-Rad CFX96 real-time PCR apparatus. SYBR Green Master mix (Bio-Rad) was used for all PCR reactions. The assay included: denaturing at 95°C for 3 min followed with 40 cycles of: 10s at 95°C and 30s at 60°C. The 2^−ΔΔCT^ method was used to calculate relative abundance of mRNA expression. Glyceraldehyde-3-phosphate dehydrogenase (GAPDH) was used as the housekeeping gene for all analyses. Primer sequences used are listed in **Table S1**.

### Histology

All tissues were fixed in 10% neutral buffered formalin and embedded in paraffin. Tissue sections (5 µm thickness) were stained with hematoxylin and eosin and mounted on slides [36,38]. Images were captured by using the cellSens Software (Olympus).

### Serum cytokine and chemokine levels

Whole blood was collected from CD-1 and B6 mice at days 2, 4, 6, 8, 10, and 12 of infection and compared with mock (PBS inoculated) controls. Serum was isolated and inactivated, as described in our previous study [36]. The Pro Mouse Cytokine 23-Plex Kit (Bio-Rad) was used to measure cytokine and chemokine levels. The Bio-Rad Bio-Plex Plate Washer and Bio-Plex machines were used for sample processing and analysis. All processes were completed following the manufacturer’s instructions.

### Statistical Analyses

Gene expression profiling data were analyzed using nSolver Software (NanoString Technologies) and presented graphically as mean ± standard error of the mean (SEM). Adjusted *p*-values were obtained using the Benjamini-Yekutieli procedure to test for significance. Data presented from survival curves, qPCR and qRT-PCR assays were analysed using GraphPad Prism software. qPCR and qRT-PCR assays are presented as mean ± SEM. Differences between control and infected groups were analyzed using Student’s t test and one-way ANOVA (parametric and non-parametric) where appropriate. Differences between survival curves were analyzed using the Log-ranked (Mantel-Cox) test. Statistically significant values are 206 denoted as \**p* < 0.05, ** *p* < 0.01, *** *p* < 0.001, and **** *p* < 0.0001, respectively.

## Results

### High susceptibility of CD-1 mice to *O. tsutsugamushi* infection

To-date, no reported studies have investigated the susceptibility of CD-1 and B6 mice in parallel via the i.v. route of *Ot* inoculation. We addressed this by performing a side-by-side infection with both mouse strains and monitoring for signs of disease. After i.v. injection with 5.6 ×10^4^ FFU of bacteria, CD-1 mice began to succumb to infection by day 9 (30%) and were completely moribund by day 12 **(Fig 1A)**. In contrast, only 10% of B6 mice succumbed on day 15 and the remaining mice (90%) recovered and survived (**Fig 1A**). Body weight loss began in CD-1 and B6 mice on days 5 and 7, respectively. Weight loss continued in CD-1 mice until host death on day 9, but weight loss ceased in B6 mice until mouse recovery on day 12 (**Fig 1B,C**). The weight loss patterns coincided closely with those of disease scores (**Fig 1D**). It is important to note that this inoculation dose (5.6 × 10^4^ FFU) was much lower than that previously used in our reports of lethal B6 models (1.25 × 10^6^ FFU) [36,38], in which tissue bacterial burdens reached a peak at day 6 [36]. As shown in **Fig 2** for infected CD-1 mice, while virtually no bacteria were detected at days 2 and 4, bacterial burdens were significantly increased at day 6 (kidneys). By day 8, bacterial burdens were significantly increased in nearly all organs (lungs, kidneys, spleen, liver), except for the brain (which reached statistical significance at day 10). The lungs and brain contained the highest vs. the lowest bacterial burdens, respectively. While spleen and liver bacterial burdens reduced to a non-significant level at day 12, other organs (lungs, kidney, and brain) maintained significantly high levels of bacteria (*p* < 0.05). These results corroborated our previous reports utilizing a lethal B6 model [36,38]. These findings indicate a systemic infection in CD-1 mice, but more importantly, a significantly higher susceptibility of CD-1 mice to *Ot* infection than B6 mice.

**Fig 1.**
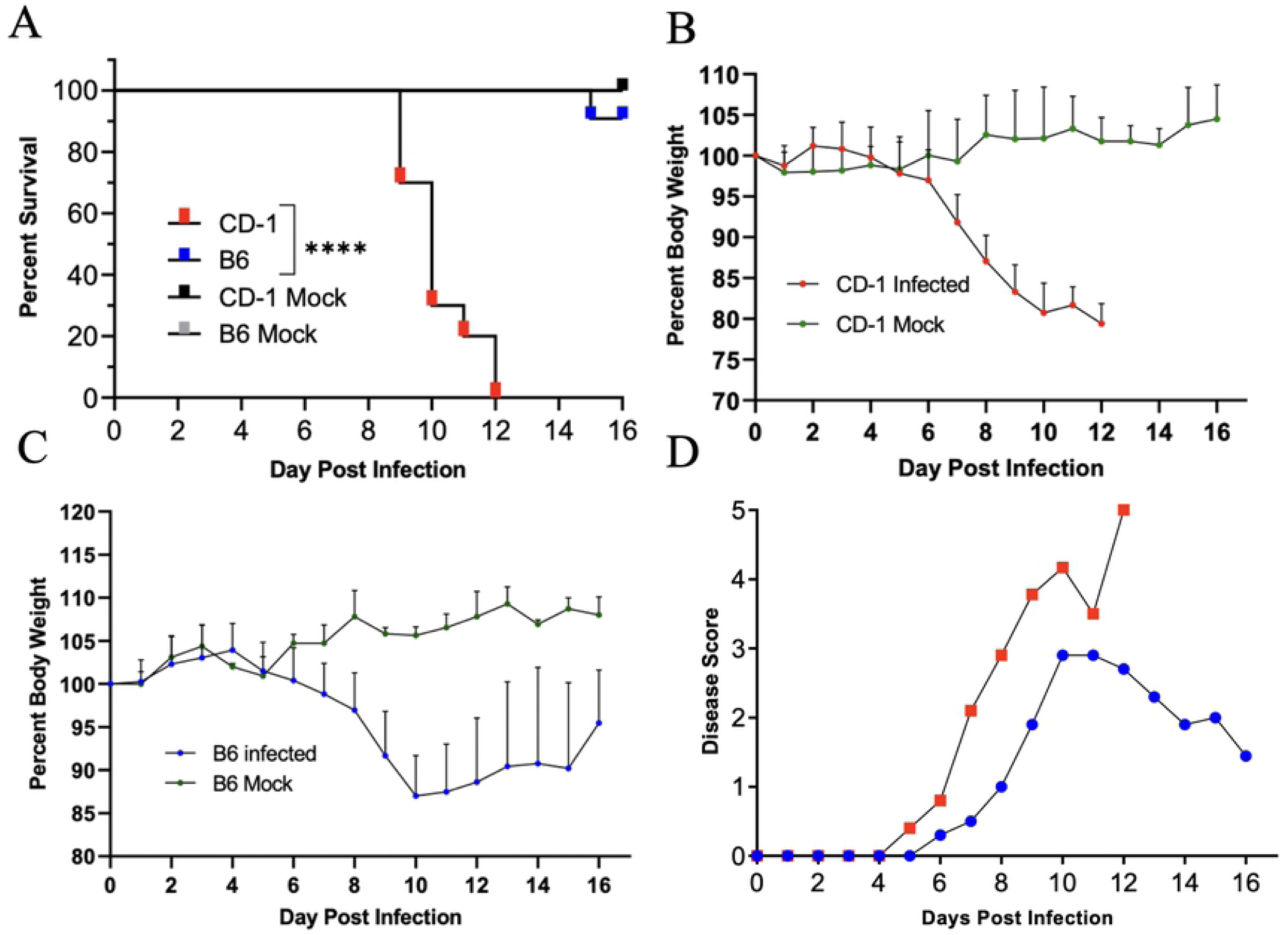
Outbred CD-1 mice were highly susceptible to *Orientia tsutsugamushi* infection. CD-1 and C57BL/6 mice were inoculated i.v. with *O. tsutsugamushi* Karp strain (5.6×10^4^ FFU, 10/group) or PBS (mock, 3/group) and monitored daily for signs of illness. (**A**) CD-1 mice began to succumb to infection on day 9, and all died by day 12. 10% of C57BL/6 died by day 15, with the rest surviving. (**B, C**) Body weight loss in CD-1 mice was rapid and progressive, in comparison to that of C57BL/6 mice. (**D**) CD-1 mice developed disease score on day 5 and had the highest score on day 12. In contrast, C57BL/6 mice started to recover at day 11. Log-ranked (Mantel Cox) test was used for statistical analysis of survival. ****, *p* < 0.0001.

**Fig 2.**
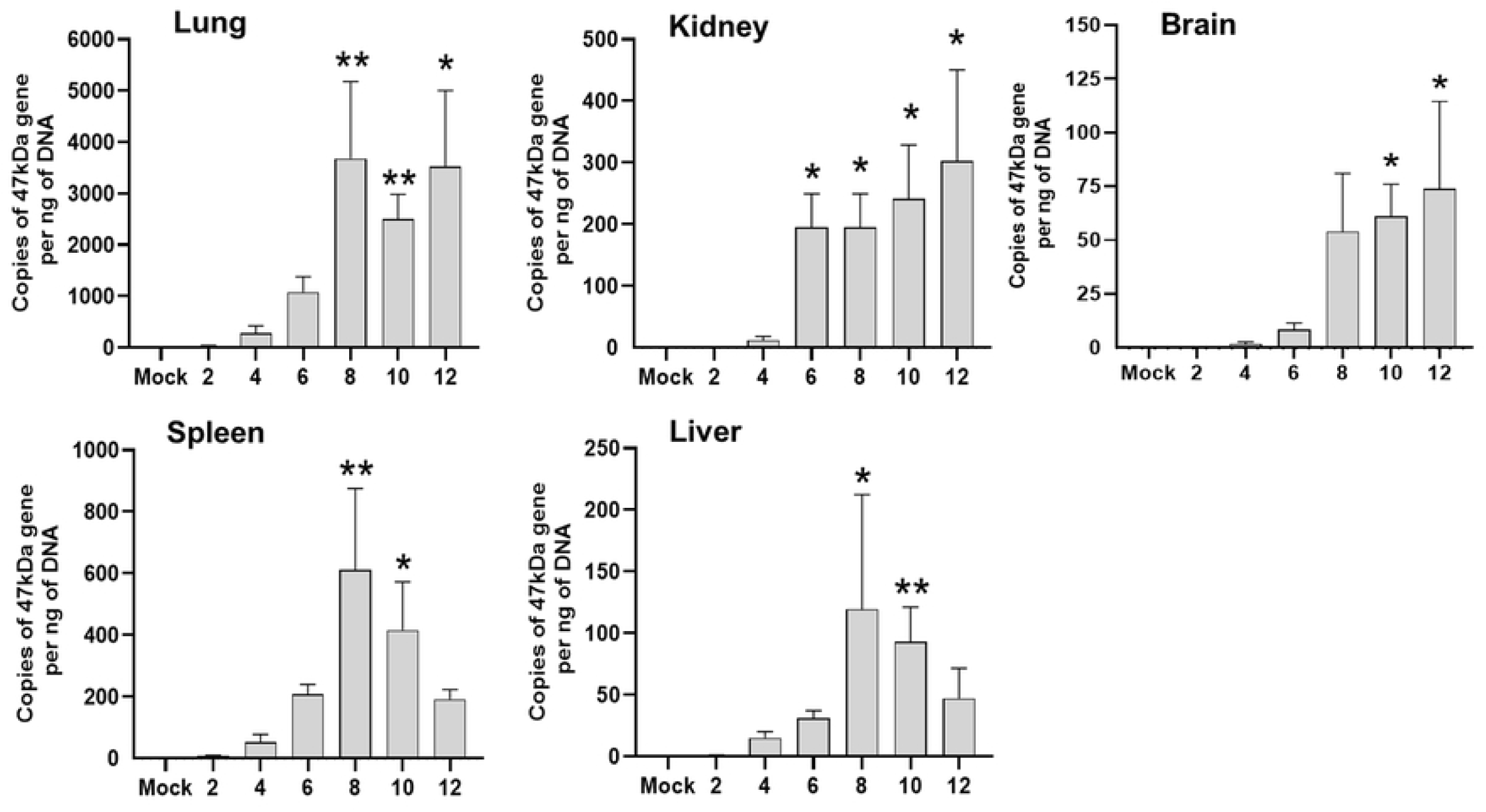
Organ bacterial burden in CD-1 mice following infection with *O. tsutsugamushi*. CD-1 mice were inoculated i.v. with *O. tsutsugamushi* Karp strain (4.32×10^4^ FFU) or PBS (mock). At indicated days post infection, organs were collected (3-4/group) for DNA extraction and bacterial burden analysis by qPCR. Data are presented as mean ± SEM. One-way ANOVA (non-parametric) was used for statistical analysis. *, *p* < 0.05; **, *p* < 0.01.

### Pathological changes and endothelial dysfunction during acute infection in CD-1 mice

Pathologic findings of human scrub typhus commonly include disseminated endothelial injury and lymphohistiocytic vasculitis, leading to interstitial pneumonitis, encephalitis, and hepatic damage [43–45]. Our established inbred mouse models mimic certain pathologic features of severe human disease [17,36,38,40,41]. As compared to mock infection (**Fig 3A**), CD-1 mice exhibited interstitial pneumonia (**Fig 3B, C**) and pulmonary alveolar edema (**Fig 3C**) indicating increased vascular permeability, consistent with our previous lethal B6 models [36,38]. CD-1 mice also showed perivascular lymphocytic infiltrates and vasculitis in the liver (**Fig S1**), as described in our previous reports [36,38], as well as multifocal mononuclear infiltrates in the cortex of the kidneys (**Fig S2**). Since *Ot* shows a tropism for replication in endothelial cells [6,43], endothelial cell activation and injury can promote recruitment, adherence, and transmigration of immune cells for immune mediated pathogen clearance and increased vascular permeability leading to edema [46]. To promote endothelial cell quiescence and vascular barrier integrity in healthy microenvironments, constitutive angiopoietin-1 (Ang1) binds the Tie2 receptor on the cell surface [47]. During infectious or inflammatory processes, Ang2 is induced and disrupts this axis, binding to Tie2 and promoting vascular barrier destabilization [48]. Therefore, the levels of *Ang1, Ang2* and *Tie2* transcripts can be used to evaluate endothelial activation/injury during infection. As shown in **Fig 4**, lung *Ang1* and *Tie2* levels were significantly decreased on day 8, while *Ang2* was insignificantly increased. The *Ang2/Ang1* ratio showed a statistically significant difference between day 8 and mock samples (**Fig 4**). This evidence of endothelial dysfunction and inflammation agrees with our previous findings in the lethal B6 mouse model [38,40,41]. Together, our study demonstrates that the CD-1 infection model mimics pathologic lesions of severe human scrub typhus in humans and inbred mouse models, offering additional value to examine biomarkers of host susceptibility and immune alterations.

**Fig 3.**
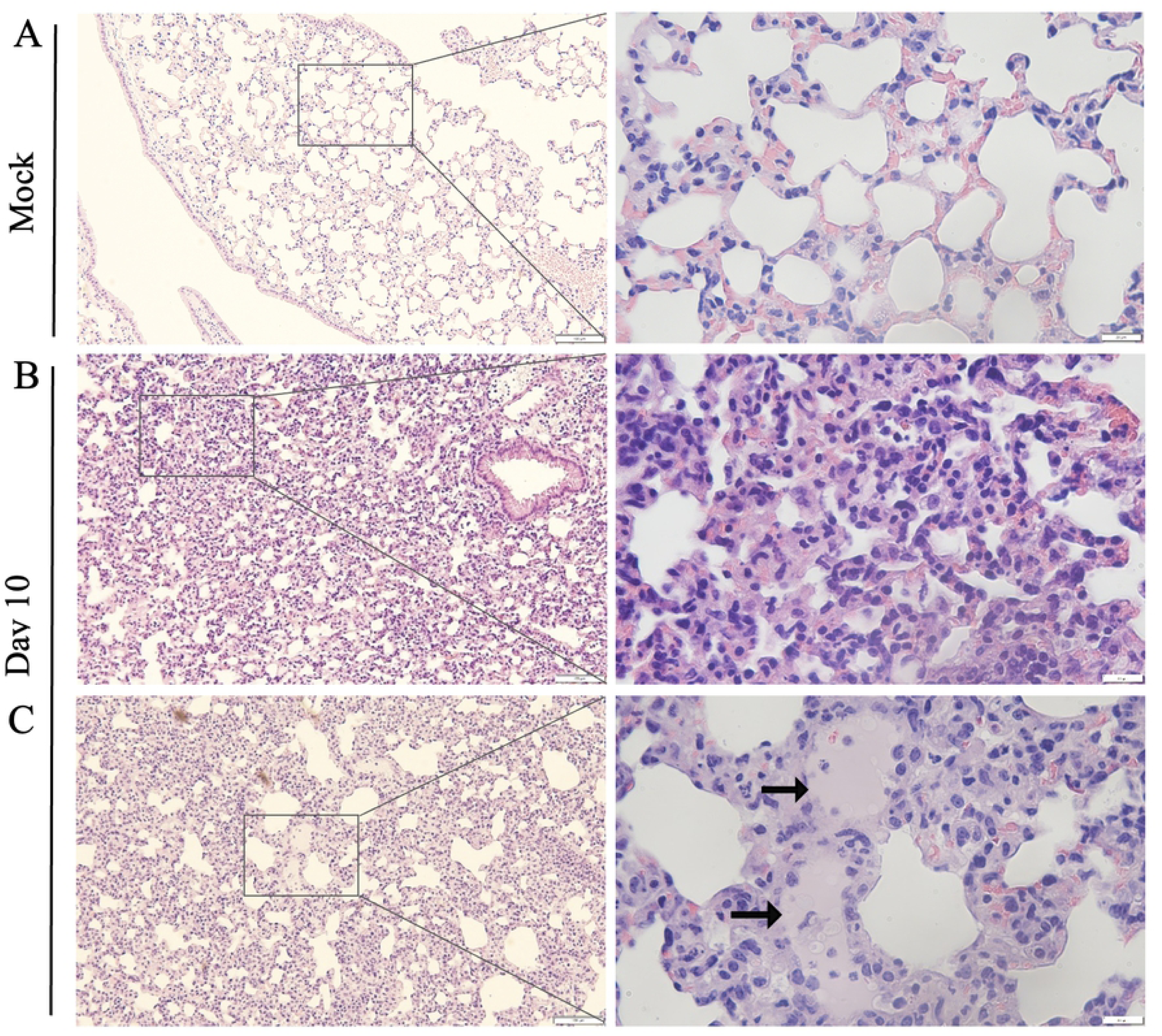
Cellular infiltration and tissue injury in *O. tsutsugamushi-*infected CD1 mice. CD-1 mice were infected as described in Figure 1. Lung tissues were collected from mock (**A**) and day 10 post-infection mice and subjected to hematoxylin and eosin staining. Cellular infiltration was present with interstitial pneumonia (**B, C**) and pulmonary edema (arrows) observed in the lung tissue of infected mice. Scale bar 100µm and 20µm (zoomed).

**Fig 4.**
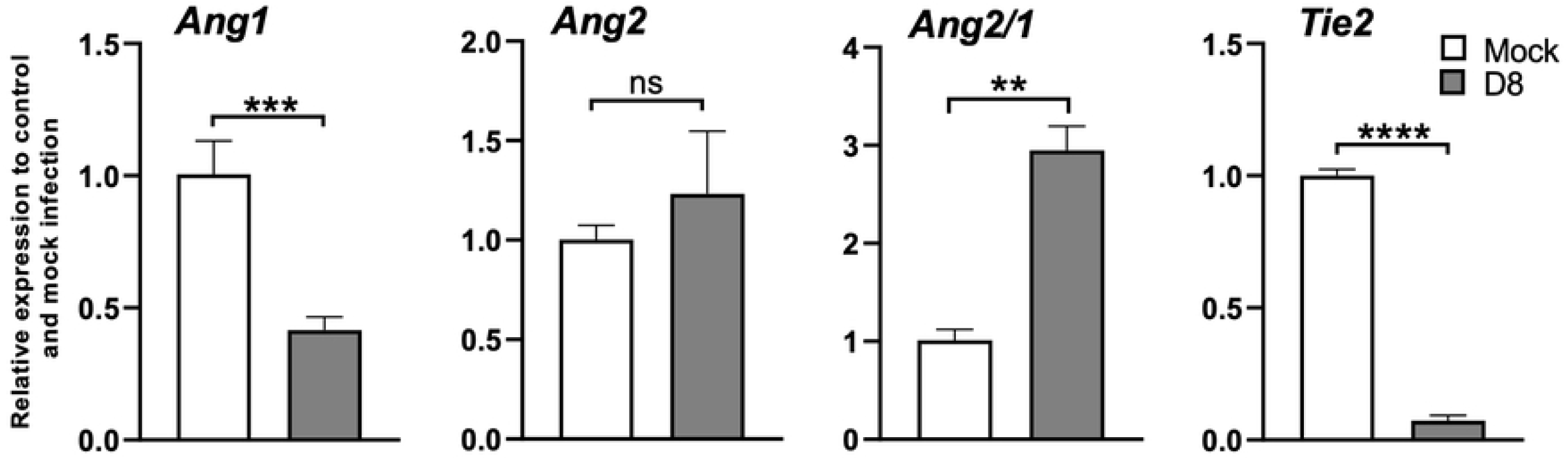
Lung endothelial dysfunction in CD-1 mice during acute infection. CD-1 mice were infected as described in Figure 2. Lung tissues were collected from mock and day 8 infected mice, and RT-qPCR assays were performed measuring *Ang1, Ang2*, and *Tie2* levels, shown as relative to GAPDH values. The Ang2/Ang1 ratio of individual samples was calculated based on RT-qPCR data and compared to mock controls. Data are presented as mean ± SEM. Students T-test was used for statistical analysis. ns = not significant; **, *p* < 0.01; ***, *p* < 0.001; ****, *p* < 286 0.0001.

### Immune profile of *O. tsutsugamushi*-infected lungs during acute infection

To gain a broad picture of immune responses in the lung during *Ot* infection in CD-1 mice, we performed pulmonary differential expression analysis on 562 immunology-related genes via NanoString. After i.v. injection with 4.32 ×10^4^ FFU of bacteria, tissues from days 4, 6, and 12 were analyzed, with mock animals serving as the baseline. Day 4 samples did not show any statistically significant changes in gene expression (see **Table S2** for complete list), which was not surprising, given the relatively low bacterial burden in tested organs (**Fig 2**, [38]), disease course (**Fig 1D**, [36]), and minimal immune responses at early stages of infection [36,38,49]. As shown in **Fig 5A** and **Table 1**, day 8 samples had 16 upregulated and 4 downregulated expressed genes of statistical significance (adj. *p-value* <0.05). At day 12, we found significantly upregulated expression of 12 genes and 1 significantly down-regulated gene (adj. *p-value* <0.05) (**Fig 5B, Table 2**). The expression kinetics of infected lungs displayed Th1-skewed immune responses during disease progression with markedly increased levels of *Cxcl9, Cxcl10, Cxcl11*, and *Gzmb* (**Fig 6A**). Other cytotoxicity-associated genes, including *Slamf7, Fasl*, and *Klrk1*, were upregulated at both days 8 and 12 (**Table 1, 2**). Meanwhile, *Il11ra1, Traf4, Illrl2*, and *Mr1* were downregulated at day 8, while *Bcap31* was downregulated at day 12 (**Tables 1, 2**). Inflammatory markers identified through our differential expression analysis were verified through qRT-PCR, confirming high expression of *Cxcl9, Cxcl10, Cxcl11, IFNγ*, and *TNFα* on day 10 (**Fig 6B**). Selectively upregulated or downregulated genes were also verified through qRT-PCR (**Fig 6C, D**), respectively. Our findings of type 1-skewed responses in *Ot* infection corroborate previous findings in the B6 models [17,38,40,42].

**Fig 5.**
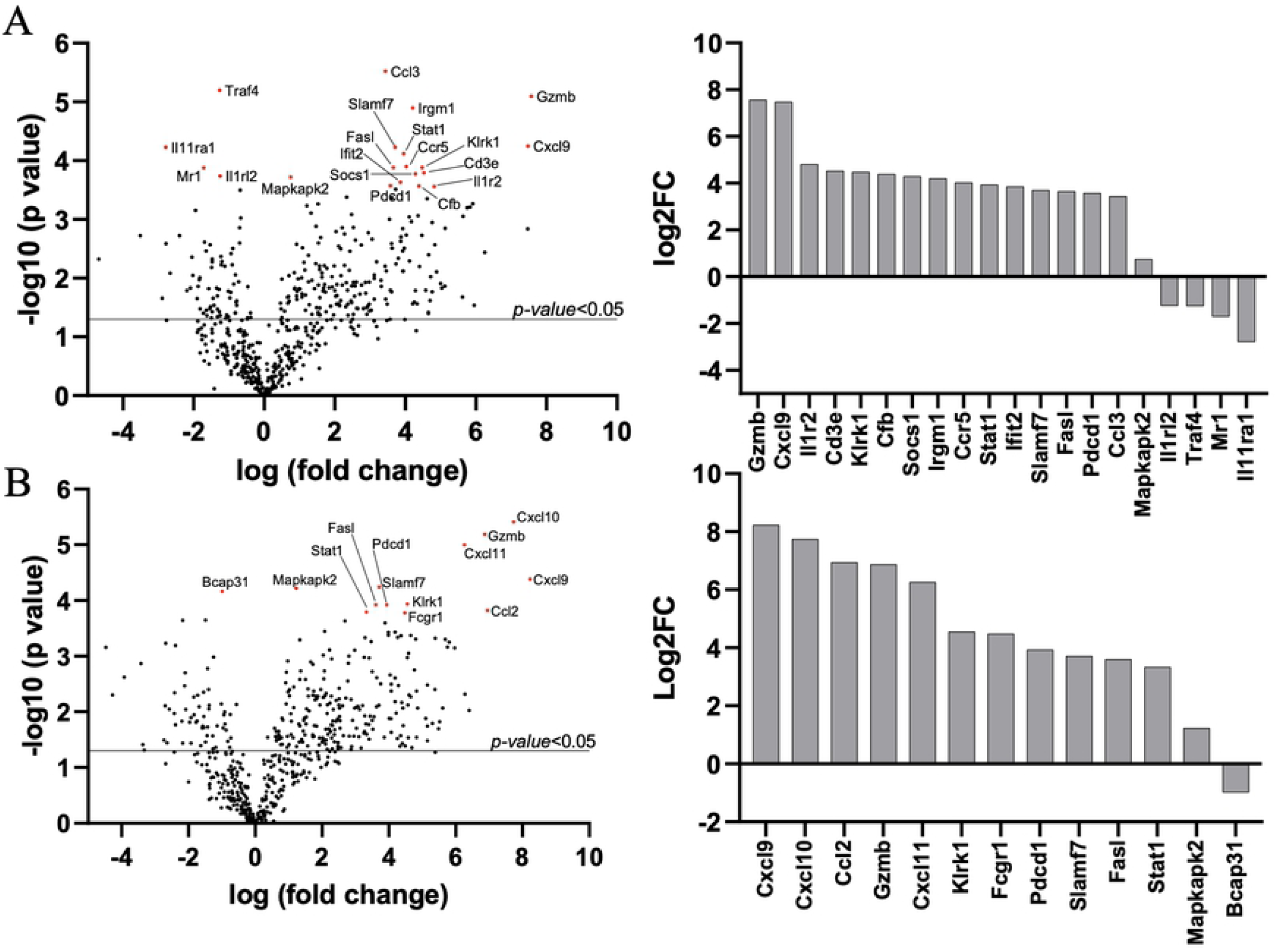
Lung transcriptomic profiling during *O. tsutsugamushi* infection. CD-1 mice were infected as described in Figure 2. Lung tissues were collected at different times (0, 4, 8, and 12 days, 3-4 mice/group) and used for total RNA extraction and Nanostring analyses. At 8 days (**A**) and 12 days (**B**) Volcano plots show significantly up-or down-regulated expression of genes (*p-value* <0.05) in black above the solid line, and significantly up-or down-regulated expression of genes with *adjusted p-value* <0.05 in red or presented in column graphs shown as Log2fold change (Log2FC).

**Table 1.**
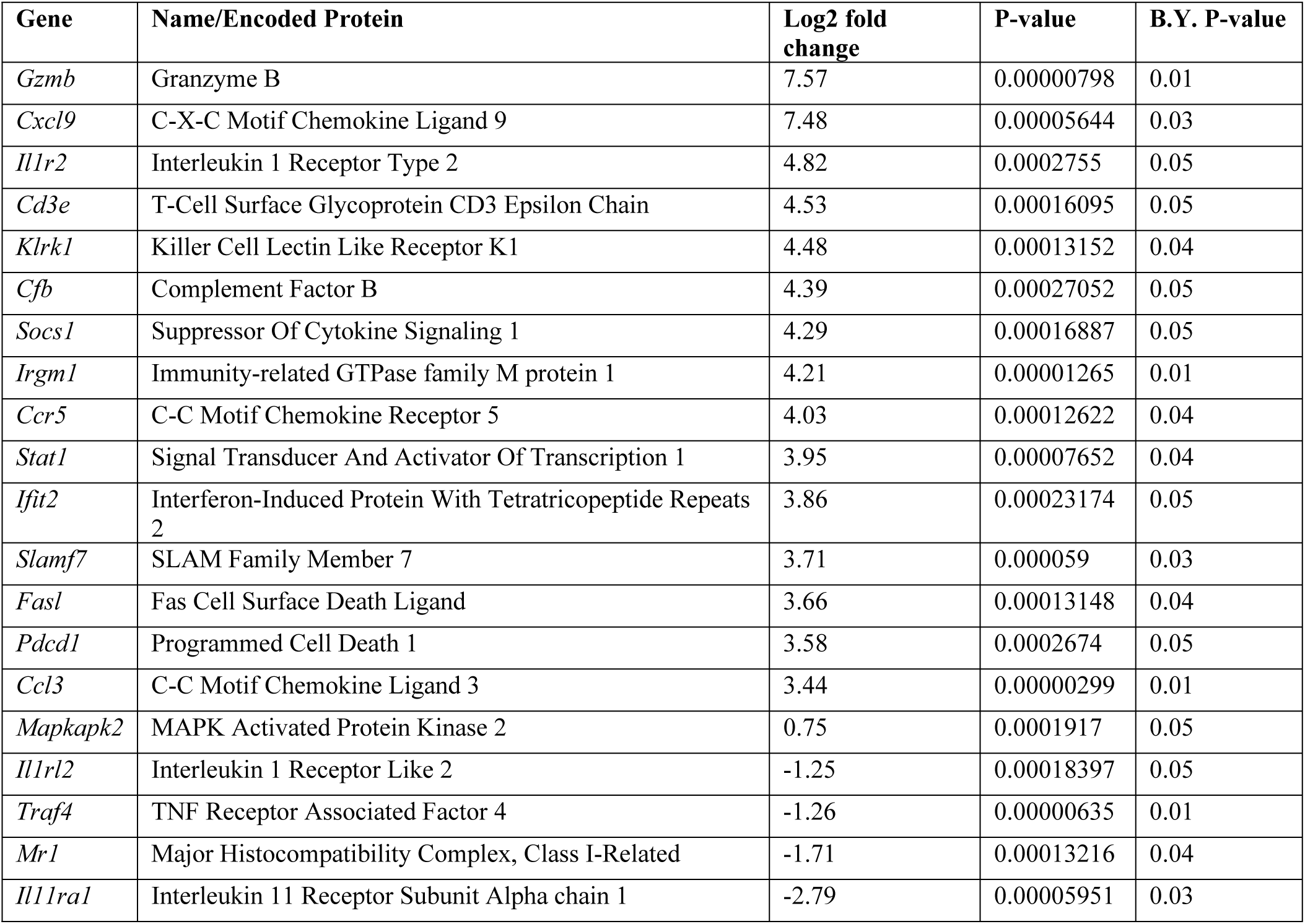
Differentially expressed genes filtered by adjusted test statistic (D8 vs. Mock)

**Table 2.** Differentially expressed genes filtered by adjusted test statistic (D12 vs. Mock)

| Gene | Name/Encoded Protein | Log2 fold change | P-value | BY P-value |
| --- | --- | --- | --- | --- |
| *Cxcl9* | C-X-C Motif Chemokine Ligand 9 | 8.23 | 0.00004141 | 0.04 |
| *Cxcl10* | C-X-C Motif Chemokine Ligand 10 | 7.74 | 0.00000386 | 0.01 |
| *Ccl2* | C-C Motif Chemokine Ligand 2 | 6.95 | 0.00015106 | 0.05 |
| *Gzmb* | Granzyme B | 6.87 | 0.00000649 | 0.01 |
| *Cxcl11* | C-X-C Motif Chemokine Ligand 10 | 6.26 | 0.0000101 | 0.01 |
| *Klrk1* | Killer Cell Lectin Like Receptor K1 | 4.55 | 0.00011555 | 0.05 |
| *Fcgr1* | Fc Fragment of IgG Receptor I | 4.48 | 0.0001675 | 0.05 |
| *Pdcd1* | Programmed Cell Death 1 | 3.94 | 0.00011924 | 0.05 |
| *Slamf7* | SLAM Family Member 7 | 3.71 | 0.00005722 | 0.04 |
| *Fasl* | Fas Cell Surface Death Ligand | 3.61 | 0.00012019 | 0.05 |
| *Stat1* | Signal Transducer and Activator of Transcription 1 | 3.33 | 0.00016157 | 0.05 |
| *Mapkapk2* | MAPK Activated Protein Kinase 2 | 1.23 | 0.00006074 | 0.04 |
| *Bcap31* | B Cell Receptor Associated Protein 31 | -0.99 | 0.00006868 | 0.04 |

**Fig 6.**
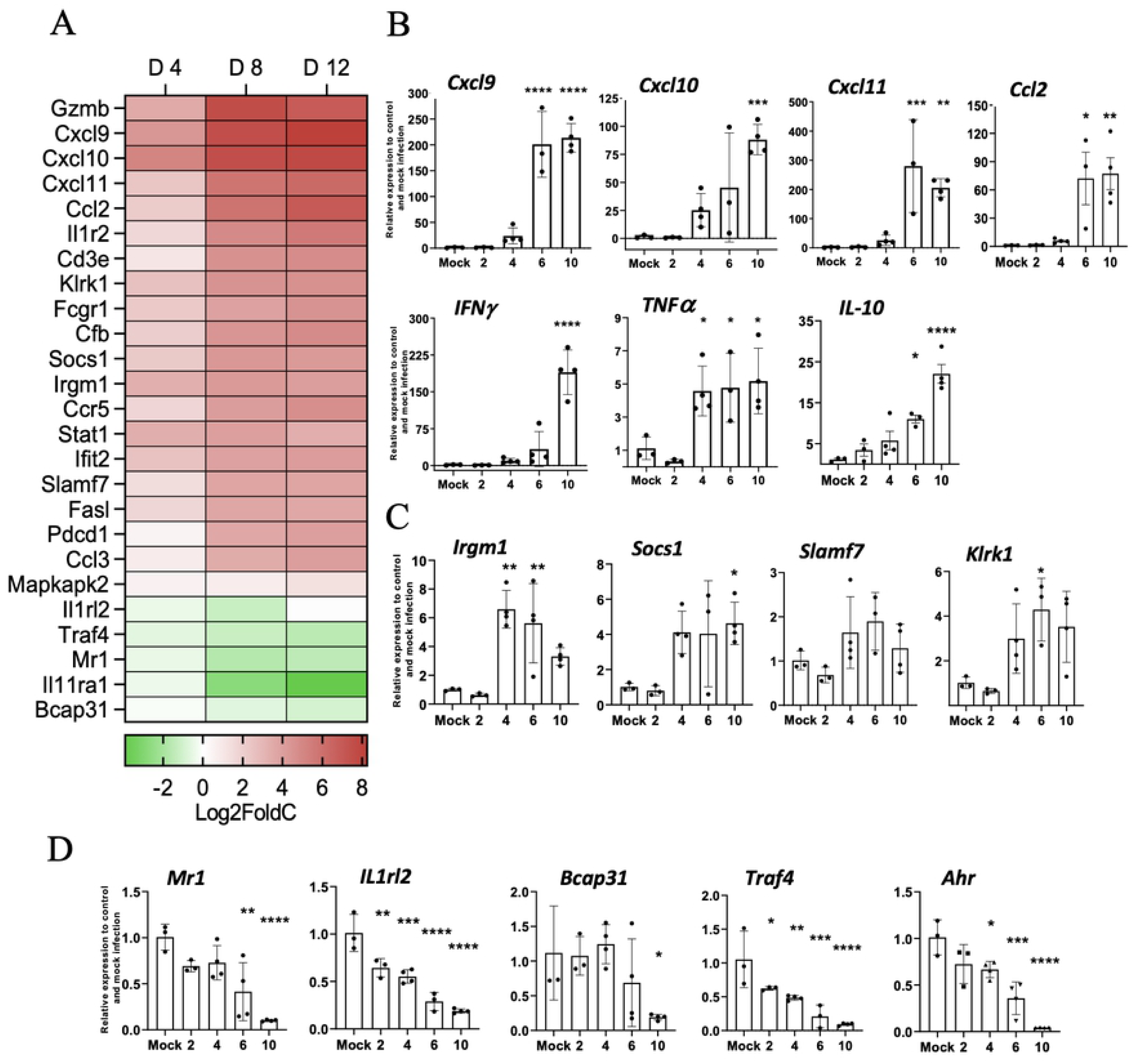
CD-1 mice displayed type 1-skewed immune profiles during acute infection. Lung transcriptomic profiling analyses were performed, as described in Figure 5. (**A**) NanoString data show Th1 and Th2 signature genes (in Log2Fold change) at indicated days post-infection as compared to mock samples. Lung qRT-PCR assays were performed and shown as relative to GAPDH values highlighting pro-inflammatory (**B**) and novel upregulated (**C**) and downregulated (**D**) markers. Data are presented as mean ± SEM. One-way ANOVA was used for statistical 328 analysis. *, *p* < 0.05; **, *p* < 0.01; ***, *p* < 0.001; ****, *p* < 0.0001.

### Other potentially important biomarkers of immune dysregulation

Considering the inherent variability associated with an outbred mouse model [27], we also parsed our gene profiling data for differentially expressed markers deemed significant (*p* < 0.05), based on unadjusted test statistics. Lung tissue expression of endothelial activation markers and receptors *VCAM-1, ICAM-1, E-selectin, Itgam, Itgal, Itga4*, and *Itgb2*, as well as endothelial-associated markers *Cd6* and *L-selectin*, were upregulated day 12, suggesting endothelial activation and damage (**Fig S3**), which were consistent with that reported in lethal B6 models [40]. Furthermore, expression of several cellular stress markers was upregulated, including *H60a, Cybb, Pdcd1, Fasl, Bid*, and *Fas*. Expression of other cytotoxicity-associated genes, including *lcos, Ctla4, Klrc1*, and *Klrd1* were upregulated (**Fig S3**). Expression of macrophage scavenger (*MARCO*, and *Msr1*) and C-type lectin receptors (*Clec4e, Clec4a4, Clec5a, Card9)* was highly upregulated (**Fig S3**). These new findings validated and expanded our recent report that Clec4e/Mincle plays an important role in sensing *Ot* and in stimulating type 1 cytokines and chemokines *in vivo* and *in vitro* [42]. In comparison, expression of Toll-like receptors was marginally or not upregulated (**Table S4**). In contrast, expression of Th2-associated markers (*IL-5, IL-13, IL-9, IL-25* and *Ahr*) was downregulated at day 12 (**Fig S3**). Together, these findings correlate with our previous studies and reveal important biomarkers and/or pathways for future studies.

### Differential cytokine/chemokine levels in sera of infected CD-1 and B6 mice

Having documented differential susceptibilities between the two strains of mice, we then analyzed cytokine and chemokine protein levels in sera by using the Bio-Plex assay. As shown in **Fig 7**, CD-1 mice, but not B6 mice, showed statistically significantly elevated levels of G-CSF at days 6 and 10, as well as significantly high levels of cytokines and chemokines at day 10 (IFNγ, TNFα, Eotaxin/CCL1, MIP-1α/CCL3, MIP-1β/CCL4, RANTES/CCL5), implying high levels of proinflammatory responses in the late stages of infection in CD-1 mice. Of note, while KC levels were highly increased in both strains of mice, the relatively resistant B6 mice had a significant KC increase at day 4 (*p* < 0.0001). In contrast, susceptible CD-1 mice showed a significant KC increase at days 6 and 10 (*p* < 0.01), revealing distinct expression kinetics of this neutrophil-recruiting chemokine. At day 10, there were statistically significant differences in levels of G-CSF, IFNγ, TNFα, Eotaxin/CCL1, MIP-1α/CCL3, MIP-1β/CCL4, RANTES/CCL5, and KC protein expression between CD-1 and B6 mice. The expression levels of IL-1α, IL-10, IL-12(p40), and MCP-1 were not significantly different. Other tested cytokines and chemokines were below detection limits, including IL-1β, IL-2, IL-3, IL-4, IL-5, IL-6, IL-9, IL-12(p70), IL-13, IL-17, and GM-CSF. Therefore, CD-1 mice generated a robust inflammatory response for cellular recruitment during the late/severe stages of the disease, in a magnitude that was much greater than similarly infected B6 mice.

**Fig 7.**
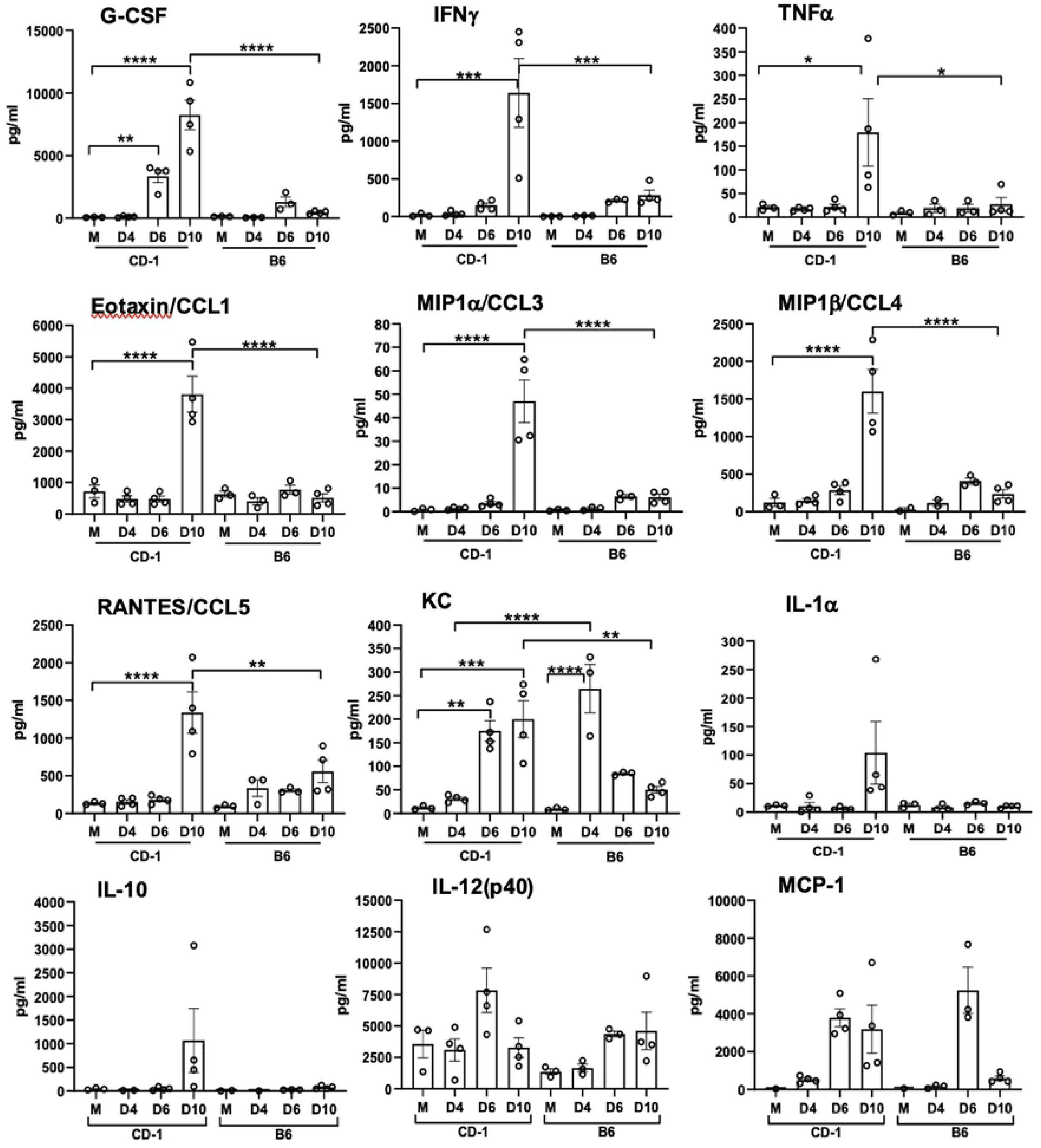
Serum protein levels of cytokines and chemokines reveal proinflammatory response in CD-1 mice versus C57BL/6 mice. CD-1 mice were infected as described in Figure 1. Whole blood was collected and serum isolated. A Bioplex assay was performed to determine cytokine and chemokine levels. Data are presented as mean ±SEM. One-way ANOVA was used for 372 statistical analysis. *, p < 0.05; **, p < 0.01; ***, p < 0.001; ****, p < 0.0001.

## Discussion

Scrub typhus is one of the most neglected tropical diseases. Obstacles to studying the biology and immunology of *Ot* bacteria have been persistent and impactful. Historically, animal model research on *Ot* has utilized the i.p. route of infection, which often causes progressive peritoneal inflammation [8,50]. Such an approach has limitations, such as accurate human-corresponding disease lesions, the disease severity produced, and translational application to humans. Here, we provide the first report of a CD-1 mouse i.v. infection model of *Ot*, as well as several lines of key findings that are important to our understanding of scrub typhus immunology and disease pathogenesis.

First, we have shown that CD-1 mice are significantly more susceptible than B6 mice to a relatively low-dose of *Ot* inoculation, judged by progressive loss of body weight, elevated disease scores, and high mortality rates (**Fig 1**). The greater susceptibility of CD-1 mice was surprising as we initially expected better infection control due to its outbred nature [23–26]. CD-1 mice did exhibit greater proinflammatory responses than B6 mice (**Fig 7**); however, host responses in CD-1 mice were either insufficient at the early stages (innate immunity) or imbalanced at the later infection stage (immune pathogenesis). Regardless of the mechanisms, our observation of mouse strain-related differences in susceptibility calls for investigation at the molecular, cellular, and organ levels. Along this line, a comparative study has indicated that CD-1-derived microglial cells demonstrated a higher scale of inflammatory responses than B6-derived microglial cells, determined by the expression levels of TNFα and IL-6 genes [51]. The pathogenic mechanisms responsible for the disparity in susceptibility between CD-1 and B6 mice warrant further investigation.

Second, it is evident this CD-1 mouse model closely resembles certain features of severe scrub typhus observed in human patients [44,45,52], as well as those seen in our previously reported lethal B6 mouse models (high dose inoculation) [36,38] regarding tissue bacterial distribution and burdens (**Fig 2**), pathological lesions (**Fig 3**), and endothelial cell dysfunction (**Fig 4**). While the highest and most sustained bacterial burden was detected in the lungs, bacterial burdens decreased to insignificant levels in the spleen and liver by day 12. Similar *Ot* distribution trends were observed in previous mouse studies employing mite bite [20,30] or i.p. injection [9], or in B6 and BALB/c mouse infection via the i.d. route [19,33]. Until the present, there were no detailed reports of CD-1 mouse infection via the i.v. route. We found that organ damage in CD-1 mice mimicked key pathological features of human disease in the lungs [52], including pulmonary edema and interstitial pneumonia with cellular infiltrates (**Fig 3**), some of which have been documented in previous reports of lethal infection in inbred mouse models [36,38,40,41]. Importantly, the pathological findings in **Fig 3** of pulmonary alveolar edema were not evident in an i.d.-inoculated B6 mouse model [17]. Liver showed perivascular lymphocyte infiltrates and vasculitis (**Fig S1**), while kidney showed mild inflammation with multifocal mononuclear interstitial infiltrates (**Fig S1**). These pathologic features have been reported in scrub typhus patients [44,45]. Like our previous reports on lethal scrub typhus in the B6 mouse lungs and kidneys [38,40,53], the CD-1 lungs showed evidence of endothelial dysfunction (high *Ang2/Ang1* ratio, but low *Tie2*) and alveolar edema during the acute disease phase (**Fig 3, 4**). This finding is important, as it reemphasizes that reduced Tie2 levels and elevated Ang1/Ang2 ratios are hallmarks of vascular dysfunction during severe stages of scrub typhus in outbred and inbred mouse strains, as well as in humans [40]. Given its broader genetic diversity compared with inbred mouse strains, *Ot*-infected CD-1 mice may serve as an excellent model for scrub typhus immunology research.

Third, we have shown that lethally infected CD-1 mice exhibit similar type 1/CD8-skewed immune profiles as in lethally infected B6 mice [38,49], and in BALB/C mice [35,37]. Our transcriptomic analysis revealed a Th1-skewed immune response during the acute phase of disease (**Fig 5**), which was further supported by significantly higher serum protein levels of IFNγ, G-CSF, IL-1α, and TNFα on day 10 (**Fig 7**). This finding was also surprising to us, as we expected a more balanced immune response due to the heterogenous nature of this mouse genetics. By providing characterization at the transcriptional and protein level of an infected outbred model, we have added additional credence to the prevalent inflammatory profile evident during lethal *Ot* infection, which helps dispel concerns that this profile is a trait solely of inbred murine models.

Our transcriptomic analysis has revealed several key biomarkers in the context of *Ot* infection. For example, the cytotoxicity gene *Gzmb* was highly upregulated at both days 8 and 12 (**Fig 5**). It is known that scrub typhus patients show high serum levels of granzymes A and B during the acute disease phase [54–58], which indicate increased cytotoxic activities and acute tissue injury [55–57]. Other cytotoxicity-associated genes revealed in our analysis (*Slamf7, Fasl*, and *Klrk1)* were also upregulated at days 8 and 12 (**Tables 1, 2**). It is possible that local production and accumulation of these cytotoxic/effector molecules contribute to endothelial cell damage and vascular dysfunction. The significantly increased CCL3, CCL4, CCL5, and CCL11 protein levels in the sera imply a robust recruitment and activation of inflammatory monocytes and macrophages at late, but not early, stages of disease (**Fig 7**). The high levels of CXCL1/2/3 protein levels in sera imply neutrophil recruitment at late stages of disease, consistent with our findings for a pathological role of neutrophils in the lethal B6 model [36]. These findings collectively suggest a strong cytotoxic response, which is an important immune response required to combat cytosolic pathogens such as *Ot* bacteria. It is known that following *Ot* infection, CD8^+^ T cells are activated in humans [59], as well as inbred mice [21], as CD8^+^ T cells are essential for survival [60] and protection against *Ot* infection [21]. Our gene profiling studies support the development of a robust cytotoxic response to *Ot* infection and lay the framework for future studies to explore the host protective versus cellular damage roles of CD8^+^ T cells during sublethal and lethal scrub typhus.

Finally, our data from CD-1 mice have provided new evidence to define biomarkers of severe scrub typhus. Our Nanostring analysis revealed several previously unidentified markers. For example, *Mr1* was one of three significantly downregulated genes during the acute phase of disease (**Fig 6A, D**). This gene encodes for an antigen-presenting protein that presents metabolites of microbial vitamin B to mucosal-associated invariant T (MAIT) cells [61]. The suppression of *Mr1* by viral infection has been shown to inhibit bacterial driven MAIT TCR-activation [62]. It is known that the *Ot* lifecycle has certain viral-like features, like closely related rickettsial bacteria [63]. A recent study of scrub typhus patients demonstrated MAIT cell activation early in infection, but MAIT cell levels were diminished at later phases of disease course [64]. *Ot* may inhibit *Mr1*, disrupt activation of MAIT cells and/or impair innate-like responses. Since the mouse and human *Mr1* genes are conserved, and mouse MAIT cells closely resemble human counterparts [65], mouse models will be beneficial for *Ot* research through characterizing the physiological and pathological roles of *Mr1* gene expression in its related cells. Moreover, we found that *Il11ra1*and *Bcap31* genes were also significantly downregulated during the acute phase of disease (**Table 1, 2**). IL-11, a cytokine in the IL-6 family, has traditionally been known to have an anti-inflammatory role, regulating fibrosis and tissue damage, while a recent study has suggested its role in epithelial dysfunction [66]. Given the downregulation of *Il11ra1* during infection (**Fig 5A**), it will be interesting to investigate whether enhanced expression of IL-11 and its related cytokines promote tissue repair. At day 12 post-infection, *Bcap31* gene was suppressed (**Fig 5B**). B-cell receptor protein 31 (BCAP31) is a ubiquitously expressed transmembrane protein with a myriad of functions and is an important membrane transport protein for the endoplasmic reticulum [67]. The suppression of *Bcap31* may be relevant to *Ot* biology, as *Ot* is known to modulate the unfolded protein response pathway through the inhibition of the endoplasmic reticulum-associated degradation pathway [68], which results in the buildup of cellular stress. Importantly, *Ot* bacteria can take advantage of this stress for its stable growth in the cytosol [68]. Therefore, active suppression of *Bcap31* may be a potential mechanism for *Ot* to promote inflammation and immune dysregulation during its establishment and replication within the host cells.

Our study also revealed several genes whose expression levels were statistically significantly different by the unadjusted test statistic (*p* < 0.05). For example, we observed that Th2-associated markers were downregulated, including *Il-4, IL-5, Il-9, Il-13, Il-25, Ccrl1*, and *Ahr* (**Fig S3**), like lethally infected B6 mice [42]. Whether such downregulation was due to the potent Th1-skewed proinflammatory responses, or to *Ot*-mediated active repression warrants further investigation.

While this is a pilot study mostly focusing on transcriptomic profiles in the lungs, it offers a broad view of the immune responses during severe scrub typhus. There are several caveats to this study, some of which will be addressed in future studies. For example, the increased sample size for CD-1 mice, especially at late/lethal stages of the disease, would greatly enhance the statistical power in our profiling analysis. The inclusion of protein-based analysis at the tissue level (immunofluorescent staining) and/or at the single-cell level (flow cytometry) would validate and expand the transcriptomic profiling data. While this study and our previous reports suggest that *Ot* infection drives a pro-inflammatory environment, mechanistic studies are needed to show how this bacterium modulates the immune response. The use of primary cell cultures would offer needed evidence to corroborate our findings of identified novel markers. Further studies will also be needed to reveal the molecular basis underlying mouse strain-specific differences in host susceptibility and immune responses following exposure to different *Ot* inoculation doses.

In summary, we have shown that CD-1 mice are more susceptible to *Ot* infection than B6 mice, showing pathologic features that mimic previous murine models and scrub typhus patients. Our results highlight the value of CD-1 mice as a good preclinical model of scrub typhus for defining host immune responses and dysregulation during severe infection. As the field of *Ot* research advances, the addition of an outbred murine model that mimics disease course, pathology, and immune response will be a vital tool for cost-effective studies to explore therapy and vaccines. Compared with inbred strains of mice, this outbred model provides a more genuine reflection of the genetically heterogeneous nature of the scrub typhus patient population. Our findings of Th1/CD8-skewed, but Th2-repressed, immunologic responses may help explain the increased susceptibility of CD-1 mice and pathogenesis of severe scrub typhus. This study will open new avenues for future mechanistic and translational studies by taking advantage of the genetic diversity in CD-1 mice to dissect the interplay between *Ot* and the host immune system.

## Acknowledgement

We thank the Baylor College of Medicine Advanced Technology Cores for performing the NanoString assays, as well as the Anatomic Pathology Laboratory at UTMB, for processing the H&E slides. We also thank Casey Gonzales and Florence Onyoni for their help in processing mouse samples during this study, as well as Dr. Keer Sun for providing helpful suggestions during the manuscript preparation.

## Supporting Information

**Supplementary Figure 1. Cellular infiltration and tissue injury in *O. tsutsugamushi-*infected CD-1 mice.** CD-1 mice were infected as described in Figure 1. Liver tissues were collected at days 0 and 10 post-infection and subjected to hematoxylin and eosin staining. Perivascular lymphocytic inflitrates and vasculitis (asterisk) were observed in the liver tissue of infected mice. Scale bar 100um and 20um (zoomed).

**Supplementary Figure 2. Cellular infiltration and tissue injury in *O. tsutsugamushi-*infected CD-1 mice.** CD-1 mice were infected as described in Figure 1. Kidney tissues were collected at days 0 and 10 post-infection and subjected to hematoxylin and eosin staining. Foci of interstitial mononuclear inflammation (dagger) were observed in the cortex of the kidney of infected mice. Scale bar 100um and 20um (zoomed).

**Supplementary Figure 3. Important biomarkers of immune dysregulation of CD-1 mice during acute infection.**1 Lung transcriptomic profiling analyses were performed, as described in Figure 5. NanoString data show differentially increased or decreased gene expression (in Log2Fold change) of distinct cellular and immune responses at indicated days post-infection as compared to mock samples.

**Supplementary Table 1.** Primers used for bacterial and mRNA quantification via qPCR and RT-qPCR, respectively.

**Supplementary Table 2.** Complete gene list from Nanostring analysis of CD-1 mice lungs on Day 4.

**Supplementary Table 3.** Complete gene list from Nanostring analysis of CD-1 mice lungs on Day 8.

**Supplementary Table 4.** Complete gene list from Nanostring analysis of CD-1 mice lungs on Day 12.

